# Structural and biochemical analysis of ATPase activity and EsxAB substrate binding of *M. tuberculosis* EccCb1 enzyme

**DOI:** 10.1101/2021.05.31.446396

**Authors:** Arkita Bandyopadyay, Ajay K. Saxena

**Author notes:** Corresponding author: Ajay K. Saxena, Rm-403/440, Structural biology lab, School of Life Sciences, Jawaharlal Nehru University, New Delhi-67, India.

## Abstract

The EccC enzyme of *M. tuberculosis* ESX-1 system is a promising target for antivirulence drug development. The EccC enzyme comprises two polypeptides (i) EccCa1, a membrane bound enzyme having two ATPase domains D2 & D3 (ii) cytosolic EccCb, which contains two ATPase domains. In current study, we have analyzed the low-resolution structure of EccCb1, performed ATPase activity and EsxAB substrate binding analysis. The EccCb1 enzyme eluted as oligomer from size exclusion column and small angle X-ray scattering analysis revealed the double hexameric structure in solution. The EccCb1 enzyme showed catalytic efficiency (*k_cat_/K_M_*)∼ 0.020±0.005 μM^-1^ min^-1^, however ∼ 3.7 fold lower than its D2 and ∼1.7 fold lower than D3 domains respectively. The D2 and D3 domains exhibited the ATPase activity and mutation of residues involved in ATP+Mg^2+^ binding have yielded 56-94% reduction in catalytic efficiency for both D2 and D3 domains. The EccCb1 binds the EsxAB substrate with *K_D_* ∼ 11.4±3.4 nM via specific groove located at C-terminal region of D3 domain. ATP binding to EccCb1 enhanced the EsxAB substrate binding by ∼ 3 fold, indicating ATPase energy involvement in EsxAB substrate translocation. We modeled the dodecameric EccCb1+EsxAB+ ATP+Mg^2+^ complex, which showed the binding pockets involved in ATP+Mg^2+^ and EsxAB substrate binding. The enzyme dynamics involved in ATP+Mg^2+^ and EsxAB substrate recognition were identified and showed the enhanced stability of EccCb1 enzyme as a result of ligand binding. Overall, our structural and biochemical analysis showed the low-resolution structure and mechanism involved in ATPase activity and EsxAB substrate binding and dynamics involved in EsxAB substrate and ATP+Mg^2+^ recognition. Overall, our structural and biochemical data on EccCb1 will contribute significantly in development of antivirulence inhibitors, which will prevent virulence factor secretion by *M. tuberculosis* ESX-1 system.

## Introduction

*M. tuberculosis* ESX-1 system (early secretory antigenic target 6 system-1) is involved in virulence factor secretion into host cell, which help mycobacterial survival in the macrophage. Loss of ESX-1 system in mycobacteria leads to genetic difference between virulent and live attenuated strains [1]. *M. tuberculosis* type VII secretion system (T7SS, ∼1.5 MDa protein complex) contains five different ESX 1 to 5 systems and involved in various physiological functions, including virulence factor secretion [2]. The function of ESX-2 and ESX-4 secretion systems are currently unknown. Recently, structure of ESX-3 secretion system was determined [3–4], which identified an additional domain in EccC enzyme, following by transmembrane helices, a stalk domain and fourth ATPase (DUF) domain. The structure of ESX-5 secretion system was also determined at 13 Å resolution using electron microscopy technique [5]. It indicates the architecture of type VII secretion system is quite different than other ESX secretion systems.

The *M. tuberculosis* EccC enzyme belongs to ESX-1 secretion system and contains (i) a cytosolic EccCb1 enzyme (Mw∼ 64.5 kDa, 591 residues) and (ii) a membrane anchored EccCa1 enzyme and belong to FTSK/SpoIIIE family of mechanoenzymes involved in DNA translocation during bacterial cell division and activator of XerCD site specific recombination [6, 7]. The EccCa1 enzyme contains two transmembrane helices, two cytosolic ATPase domain and forms complex with EccCb1enzyme and export EsxAB substrate out of the cell [2, 6]. The EccCb1 enzyme contains two P-loop NTPase domains, as observed in ACSE (Additional Strand Conserved Glutamate) family of proteins e. g. Ftsk, TrwB and TrwK [8]. The ASCE family of enzymes harbor the residues, which enters into active site of neighboring domain upon multimerization [9]. The *M. tuberculosis* EccC enzyme consists of two polypeptides, while single polypeptide is observed for *T. curvata* EccC enzyme [10] and *G. thermodentrificans* EssC enzyme [11]. In *T. curvata* EccC enzyme, linker between ATPase1 and ATPase2 domains was found quite similar to the linker between ATPase2 and ATPase3 domains and play key role in regulation of ATPase1 domain activity.

*M. tuberculosis* EsxAB substrate belongs to WxG superfamily of proteins [12–13]. The WxG family of proteins form characteristic homo or heterodimer, in which one monomer contains the WxG motif and another monomer contains the YxxxD/E motif in its export arm. The export arm of EsxAB substrate binds to EccCb1 enzyme for its secretion out of the cell [14]. The ESX-1 substrates secrete in codependent manner, as *M. tuberculosis* strains lacking the EsxAB substrate were unable to secrete other virulence factors out of the cell [15].

Recently, crystal structure of C-terminal D3 domain of *M. tuberculosis* EccCb1 enzyme in complex with peptide from EsxB substrate was determined at 2.2-Å resolution [16]. The structure analysis showed the nature and location of substrate binding pocket of EccCb1 enzyme. In another study, crystal structure of ATPase domain of EssC enzyme from *S. aureus* USA300_FPR3757 was determined at 1.7-Å resolution [17], which also elucidated the structure of substrate binding pocket of the enzyme.

To understand the structure and mechanism of EccCb1 enzyme, we have determined the low-resolution structure of EccCb1 using small angle X-ray scattering technique. The ATPase activity of full length Rv3871 and its D2 and D3 domains were determined and performed comparative analysis. The EsxAB substrate binding to EccCb1was determined and enzyme dynamics involved in ATP+Mg^2+^ and EsxAB substrate recognition were determined using dynamics simulation technique. Our structural and biochemical data on EccCb1 can be targeted for anti-virulence inhibitors development, which will prevent virulence factor secretion by *M. tuberculosis* ESX-1 system.

## Results

### Full length EccCb1 forms dodecamer in solution and exhibited specific ATPase activity

The schematic view of full-length EccCb1 enzyme (65 kDa, 591 residues) is shown in **Fig. 1A**. The EccCb1 consists of two polypeptides, D2 domain (35-316 residues) and D3 domain (349-580 residues), which belong to P- type NTPase family. The ATP binding motif of D2 domain (^84^GAPQTGKS^91^, cyan) and D3 domain (^376^GAAKSGKT^383^, brown) are shown in **Fig. 1A**. Full length EccCb1 were purified, however eluted as oligomer on Superdex200^TM^ (16/60) column, identified based on molecular mass standard **(Fig. 1B, inset).** The eluted protein from column showed purity more than 98% and single band on SDS-PAGE analysis (**Fig. 1B, inset**). Small angle X-ray scattering analysis on Rv3871 showed double hexameric structure (Mw ∼ 780 kDa) in solution.

**Fig. 1.**
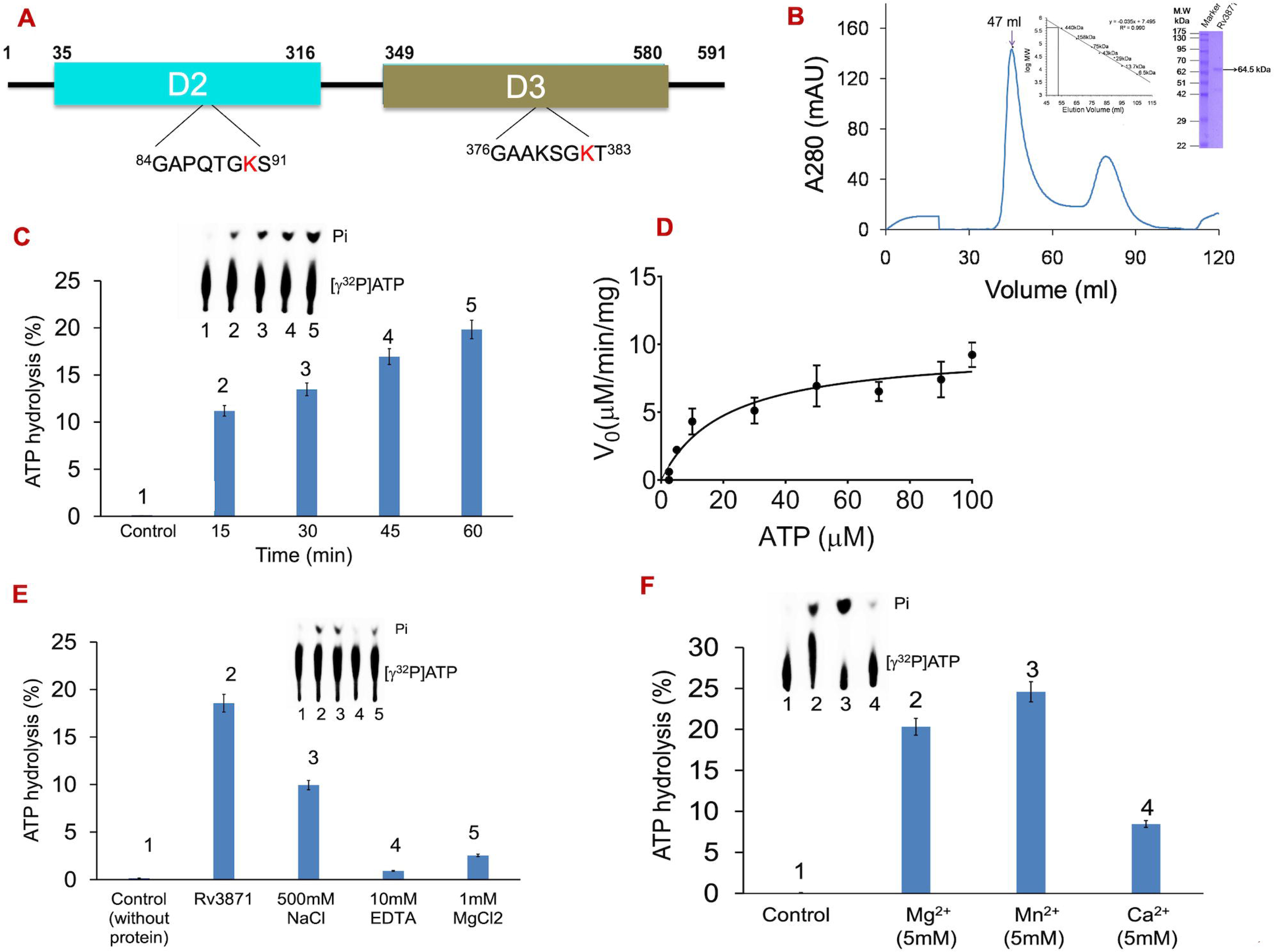
Purification and ATPase activity of EccCb1 enzyme. **A,** Schematic diagram of full length EccCb1 (1-591 residues), consists of D2 domain (35-316 residues, blue) and D3 domain (349-580 residues, cyan). The 84-91 residues of D2 domain and 376-383 residues of D3 domain are involved in ATPase activity and highly conserved in Ftsk/SpoEIII family of enzymes. **B,** Elution profile of cleaved EccCb1 from Superdex^TM^ 200(16/60) column. The protein eluted as oligomer (based on molecular mass standard, *inset*) and appeared as single band on SDS-PAGE *(inset).* **C,** Time dependent ATP hydrolysis of EccCb1 enzyme. I*nset* shows the radioactivity profile of the ATP hydrolysis. **D,** ATPase activity of EccCb1 enzyme using malachite green assay and *K_m_, k_cat_* and *V_max_* parameters were obtained using Graph Pad software. **E,** Percentage of ATP hydrolysis by EccCb1 in presence of 10 mM EDTA, 500 mM NaCl and 1 mM MgCl_2_. *Inset* shows the radioactivity profile of ATP hydrolysis. **F,** Percentage of ATP hydrolysis by EccCb1 in presence of different divalent cations. *Inset* shows the radioactivity profile of ATP hydrolysis

[γ-^32^P] radioactivity assay was performed to analyze the ATPase activity of EccCb1 enzyme **(Fig. 1C).** In radioactivity assay, release of free Phosphate was monitored gradually after every 15 min and showed maximum ATP hydrolysis (20%) at 60 min (**Fig. 1C**). Three measurements were taken for each reading and reaction buffer containing no EccCb1 taken as negative control. Calorimetric assay showed the ATPase activity of purified EccCb1 and following parameters e. g. *V_max_* ∼ 9.28 μM min^-1^ mg^-1^, *K_m_* ∼ 0.02±0.01 mM and *k_cat_/K_m_* ∼ 0.020±0.005 μM^-1^ min^-1^ were obtained using Michaelis-Menten plot **(Fig. 1D)** and Graph Pad Prism 6.0 software [18].

To examine the effect of various parameters e. g. oligomerization, EDTA, MgCl_2_ on EccCb1 ATPase activity, [γ-^32^P] radioactivity assay was performed using these compounds. In presence of 500mM NaCl in reaction buffer, the EccCb1 showed 50% decrease in ATP hydrolysis. In presence of 10mM EDTA, the EccCb1 showed no ATP hydrolysis indicating absolute requirement of Mg^2+^ for enzyme activity. Mg^2+^ binds to β- and γ- PO_4_ of ATP and neutralize the negative charge on PO_4_ group and promotes the nucleophilic attack by activated water for ATP hydrolysis. In presence of 1 mM MgCl_2_, EccCb1 showed only 2% ATP hydrolysis compared to wild type enzyme **(Fig. 1E)**.

To analyze the effect of various ions on percentage of ATP hydrolysis, [γ-^32^P] radioactivity assay was performed and (%) of ATP hydrolysis was calculated. 25% ATP hydrolysis was observed with 5mM Mn^2+^ 8% with 5mM Ca^2+^ and 20% with 5mM Mg^2+^ **(Fig. 1F)**. These data showed that Mn^2+^ ion (radii ∼ 0.65Å) fit better into ATPase pocket than Mg^2+^ ion (ionic radii ∼ 0.46Å), which resulted in higher ATPase activity of EccCb1 enzyme. However, Ca^2+^ (ionic radii ∼ 1.11 Å) is higher than Mg^2+^ (radii ∼ 0.46Å) and may not be fitting optimally into ATPase pocket, which result in lower ATPase activity of EccCb1 enzyme.

### Small angle X-ray scattering analysis yielded the double hexametric ring structure of EccCb1 in solution

To determine the low-resolution structure of EccCb1, small angle X-ray scattering experiment was performed. The SAXS profile obtained in double logarithmic mode **(Fig. 2A)** supported a monodisperse profile of EccCb1 in solution, devoid of any aggregation or interparticulate effect. Considering the monodisperse globular shape of EccCb1, linearity in the Guinier region was observed **(Fig 2A, inset).** Globular scattering profile of EccCb1 in solution was confirmed by peak close to 1.75 in normalized Kratky plot **(Fig. 2B).** The Guinier and Kratky plots analysis yielded the radius of gyration (R_g_) ∼ 6.87 nm for EccCb1. Presuming the rod-shape structure of EccCb1, the Guinier analysis showed the cross-sectional radius, R_c_ ∼ 6.77 nm. Considering wider q range of data, P(r) analysis on EccCb1 yielded the D_max_ ∼ 20.14 nm and R_g_ ∼ 6.86 nm **(Fig. 2C).** Peak-shoulder of the P(r) curve showed single lobe of scattering **(Fig. 2C).** Using dummy residue models and shape restoration using SAXS data-based constraints showed an envelope compatible to double hexameric ring structure of EccCb1 **(Fig. 2D).** The NSD value ∼ 0.883± 0.132 was observed for selected model with resolution ∼ 9.7±0.7 nm. The theoretical R_g_ and D_max_ were quite compatible to the experimental R_g_ and D_max_ values.

**Fig 2.**
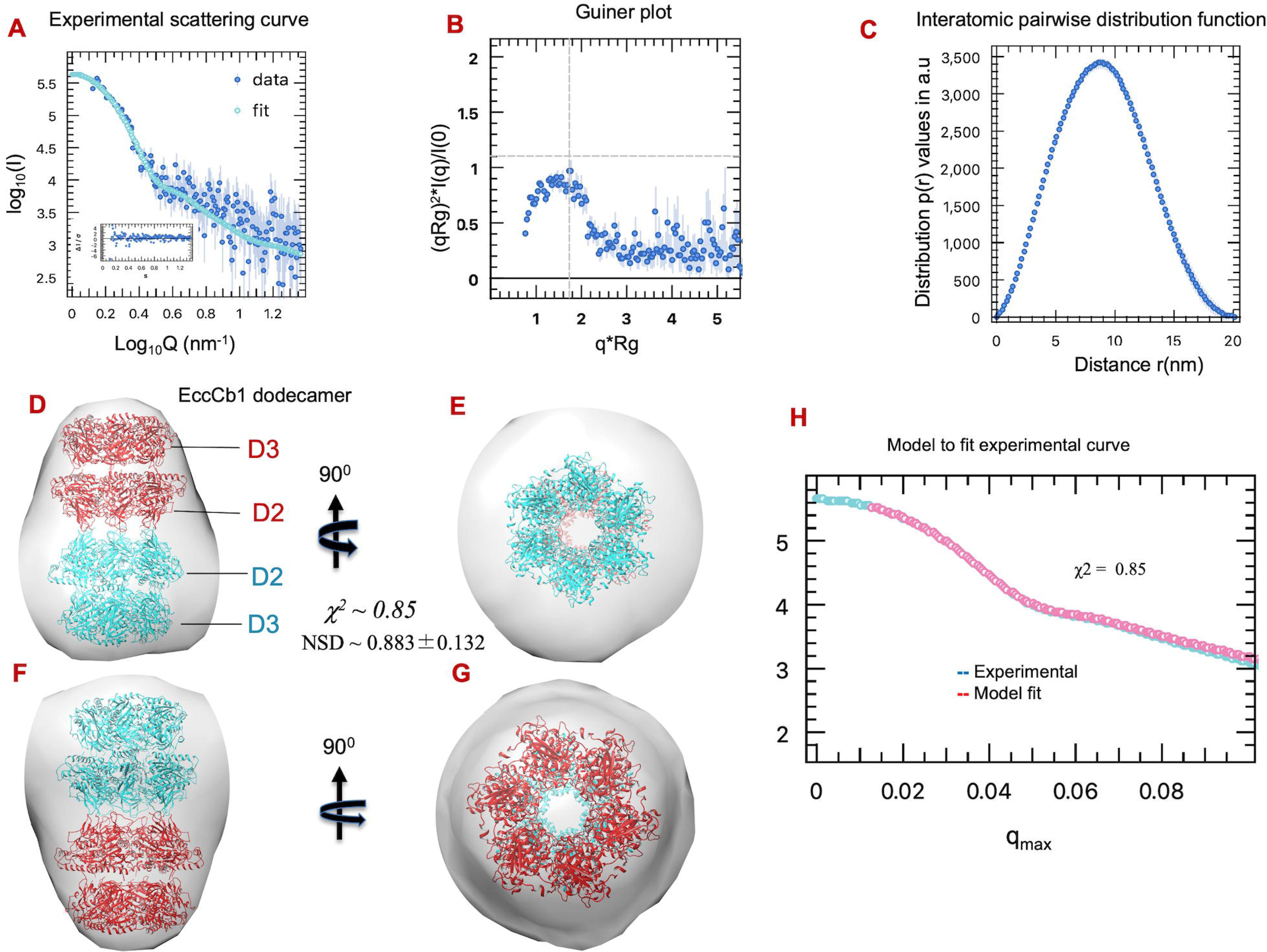
Small angle X-ray scattering analysis of EccCb1 enzyme. **A**, Experimental SAXS intensity profile of EccCb1. *Inset* shows the linear fit to Guinier region of measured data. **B,** Guinier plot in low q region showing linear agreement between experimental data (blue crosses) and Guinier equation (blue line) and further confirmed by low residuals within boundaries. **C,** Inter-atomic pairwise distribution function, P(r), calculated by GNOM program, which indicates the frequency distributions of interatomic vectors in predominant scattering species. At D_max_ ∼ 20 Å, the P(r) function approaches zero smoothly. **D-E,** 0 and 90° views of double hexameric ring of EccCb1 fitted in molecular envelope generated by *ab initio* SAXS modeling. **F-G,** 0 and 90° views of double hexameric ring of EccCb1 (180° view of **Fig. D-E**). **H,** Fitting of experimental scattering data (cyan) with theoretical scattering curve (red) obtained from 10 *ab initio* models generated by DAMMIF program. To generate envelope, DAMAVER superimposed all 10 models, averaged and filtered the final model to generate envelope, as shown in **Fig. 5D-G**. The mean normalized spatial discrepancy (NSD) of 0.883±0.132 observed for 10 aligned models.

The *ab initio* envelope of EccCb1 was obtained from SAXS data. It indicates the presence of double hexameric ring structure of EccCb1 in solution. We modeled the double hexameric ring structure of EccCb1, which fitted very well into acquired SAXS envelope **(Fig. 2D-G).** The CRYSOL program [19] was used to compare theoretical SAXS profiles from model vs experimental SAXS data, which yielded quite good fitting (χ^2^ ∼ 0.85) **(Fig. 2H).** The details of SAXS data collection and various parameters used in structure solution are given in **Table 3**.

### D2 domain forms dimer in solution and exhibited ATPase activity

The web logo diagram [20] of sequence alignment of ATPase motif of D2 domain with 80 heterologous EccC sequences is shown in **Fig. 3A**. These data showed that ^90^GKS^92^ sequence of ATPase motif is 100% conserved in all 80 EccC sequences. The D2 domain (35-316 residues, 31kD) was purified and eluted as dimer from Superdex 200^TM^ (16/60) column, as identified based on molecular mass standard **(Fig. 3B, inset).** The D2 domain showed purity more than 98% and appeared as single band on SDS-PAGE analysis (**Fig. 3B, inset).** Four site directed mutants of D2 domain were prepared and all mutants eluted as dimer like wild type D2 domain (**Fig. 3B)**.

**Fig. 3.**
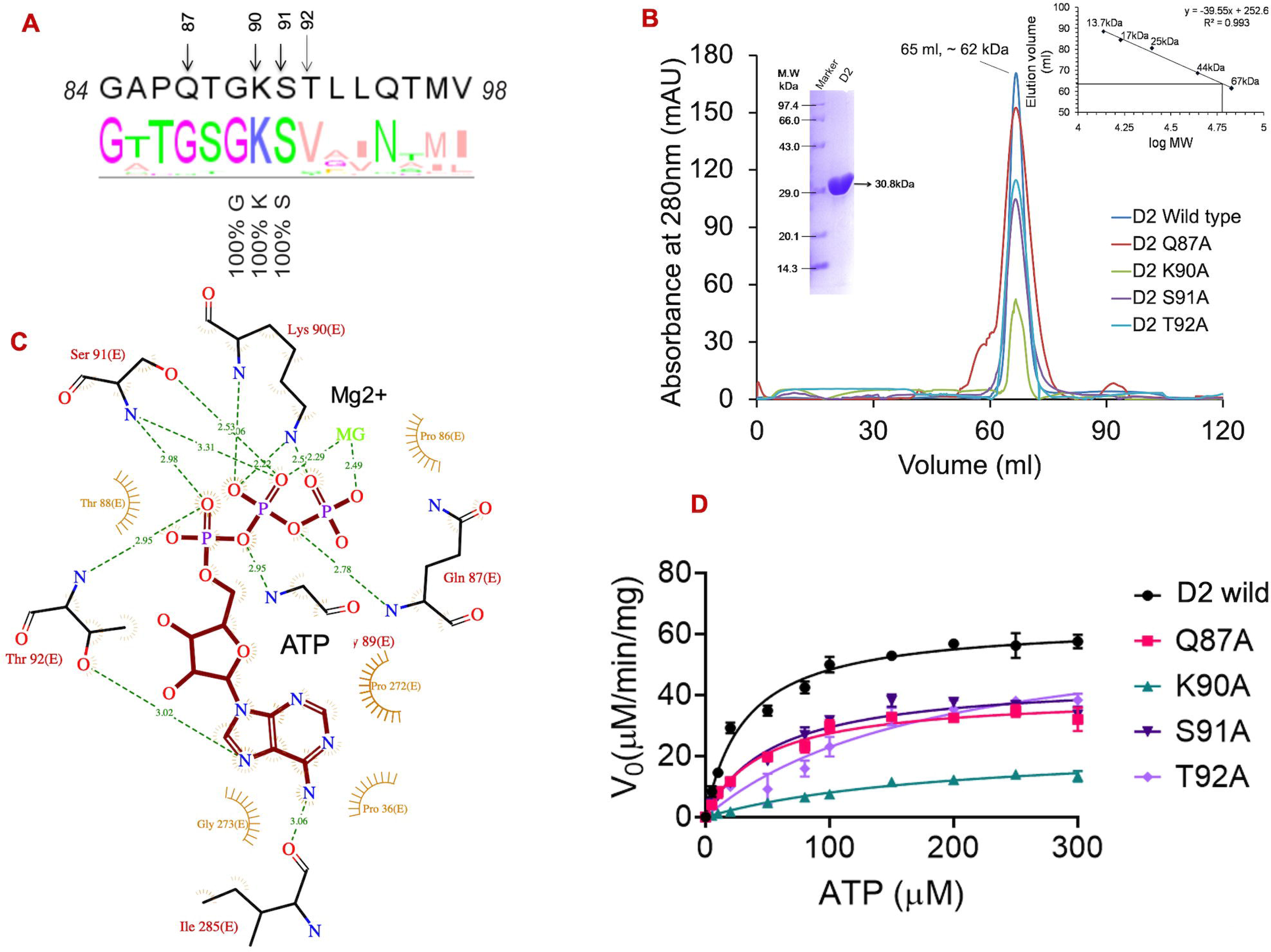
Purification and ATPase activity of wild and mutant D2 domains. **A,** Web logo diagram of ATPase motif of D2 domain aligned with Ftsk/SpoEIII motif of 80 EccC sequences. **B,** Elution profile of wild and four mutant D2 domains obtained from Superdex200^TM^ (16/60) column and eluted as dimer from size exclusion column. **C,** LigPlot analysis of D2+ATP+Mg^2+^ complex showing the interactions between ligand and D2 domain. ATP is shown in maroon, Mg^2+^ in green and all interacting residues in red. **D,** Michaelis-Menten plots showing the wild and four point mutant (Q87A, K90A, S91A, T92A) D2 domains.

ATPase activity of D2 domain was determined using calorimetric assay and following parameters (*K_m_* ∼ 0.03±005 mM, *V*_max_ ∼ 63.79 μM/min and catalytic efficiency (*K_cat_/K_m_*) ∼ 0.070±0.020 μM^-1^ min^-1^) were obtained using Michaelis-Menten plot and nonlinear regression analysis using Prism 6.0.2 GraphPad Software **(Fig. 3D).** We modeled the D2 domain+ATP+Mg^2+^ complex based on PDB-4N1A [10] structure as input and identified key residues involved in ATP and Mg^2+^ binding using LigPlot program [21] **(Fig. 3C).** The Gln87, Lys90, Ser91 and Thr92 residues of D2 domain form hydrogen bonds with γ-PO4 group of ATP and Thr92 and Ile285 residues form hydrogen bonds with adenine ring of ATP.

To understand the roles in ATPase activity, we generated the Q87A, K90A, S91A and T92A D2 mutants and analyzed the ATPase activity **(Fig. 3D). Table 1** showed the kinetic parameters observed for four D2 mutants. The Q87A mutation leads to (∼ 55.9% decrease in catalytic efficiency and ∼ 1.5-fold decrease in V_max_). The K90A mutation leads to (∼ 93.9% decrease in catalytic efficiency and ∼ 2.6-fold decrease in V_max_). The S91A mutation leads to (∼ 42.2% decrease in catalytic efficiency and ∼ 1.4-fold decrease in V_max_). The T92A mutation leads to (∼ 82.3% decrease in catalytic efficiency and quite similar V_max_) compared to wild type D2 domain. These data showed that K90A mutation critically affect the ATPase activity and leads to almost complete loss of D2 domain activity. The wild type and four D2 mutants were expressed and purified under similar conditions and showed no abnormal elution behavior on size exclusion column **(Fig. 3B).** Differences in ATPase activity of four D2 mutants were not due to defect in folding or aggregation of proteins.

**Table 1.** Kinetic parameters of full length EccCb1 and its D2 & D3 domains.

| Protein | $V_{max}$<br>$\mu\text{M}/\text{min}/\text{mg}$ | $K_m$<br>$\text{mM}$ | $k_{cat}$<br>$\text{min}^{-1}$ | $k_{cat}/K_m$<br>$\mu\text{M}^{-1}\text{min}^{-1}$ | (%) of<br>activity<br>observed |
| --- | --- | --- | --- | --- | --- |
| **Full length**<br>**EccCb1** | **9.28** | **0.02±0.01** | **0.31±0.05** | **0.020±0.005** |  |
| **D2 domain** | **63.79** | **0.03±0.01** | **2.13±0.10** | **0.070±0.020** | **100%** |
| D2-Q87A | 39.88 | 0.05±0.01 | 1.33±0.08 | 0.030±0.007 | 44.1% |
| D2-K90A | 23.92 | 0.20±0.06 | 0.80±0.13 | 0.004±0.002 | 6.1% |
| D2-S91A | 45.22 | 0.05±0.02 | 1.15±0.13 | 0.030±0.008 | 42.2% |
| D2-T92A | 65.22 | 0.19±0.10 | 2.18±0.60 | 0.010±0.006 | 17.7% |
| **D3 domain** | **52.89** | **0.06±0.03** | **1.76±0.27** | **0.030±0.008** | **100%** |
| D3- K382A | 17.49 | 0.40±0.28 | 0.60±0.10 | 0.002±0.001 | 6.6% |
| D3- T383A | 26.34 | 0.30±0.20 | 0.88±0.50 | 0.003±0.002 | 10.3% |
| D3- T384A | 41.70 | 0.40±0.10 | 1.40±1.00 | 0.004±0.001 | 12.7% |
| D3-A574G | 46.76 | 0.20±0.08 | 1.56±0.35 | 0.008±0.004 | 26.5% |
| D3- Y576A | 45.21 | 0.20±0.08 | 1.51±0.30 | 0.008±0.004 | 24.8% |

**Table 2.** SAXS data collection and *de novo* structure analysis of EccCb1

| Instruments/programs | data analysis |
| --- | --- |
| **SAXS Data Acquisition** |  |
| Instrument | SAXSpace (Anton Paar) (At CSIR-IMTECH) |
| Collimation | Line |
| Wavelength (nm) | 0.154 |
| q range (1/nm) | 0.1 – 5 |
| Exposure time | 30 minutes (Frames of three for averaging) |
| Protein Concentration | 5 mg/ml |
| Temperature | 283 K |
| **Programs** |  |
| Data Collection | SAXSDrive |
| Beam position correction | SAXSTreat |
| Buffer subtraction | SAXSQuant |
| Desmearing | SAXSQuant |
| **Data Analysis** |  |
| All parameter analysis | SAS Data Analysis (ATSAS v 3.0.1)<br>Guinier Analysis, Distance Distribution, Molecular Weight Estimation |
| Shape Restoration | Dummy residue model<br>10 models with online DAMMIF<br>DAMCLUST for clustering into families<br>DAMAVER for Averaging |
| Inertial Alignment | SUPALM (Pymol), Supcomb (Command Line) |
| Theoretical SAXS calculation | CRY SOL |
| 3D Graphics | PyMol, UCSF Chimera |
| Guinier based $R_g$, $M_w$ | 6.87nm, 717.6 kDa |
| P(r) based $D_{max}$, $R_g$ | 20.14 nm, 6.86 nm, |
| NSD of models | 0.883±0.132 |
| Resolution of ensemble | 9.7±0.7 nm |
| *Calculated $\chi^2$ of theoretical SAXS vs. Experimental Data* |  |
| EccCb1 model | 0.85 |
| Molmap Generation Scale | 20 for DAMFILT and DAMAVER |

### D3 domain forms dimer in solution and exhibited ATPase activity

Web logo diagram of sequence alignment of ATPase motif of D3 domain with 80 heterologous EccC sequences is shown in **Fig. 4A**. These data showed that Gly382 and Lys383 residues of D3 domain were 86% and 84% conserved in all EccC sequences. The D3 domain eluted as dimer (Mw∼ 50 kDa) from Superdex200^TM^ (16/60) column, as identified based on molecular mass standard **(Fig. 4B, inset).** Purified D3 domain showed purity more than 98% and appeared as single band on SDS-PAGE analysis **(Fig. 4B, inset).**

**Fig. 4.**
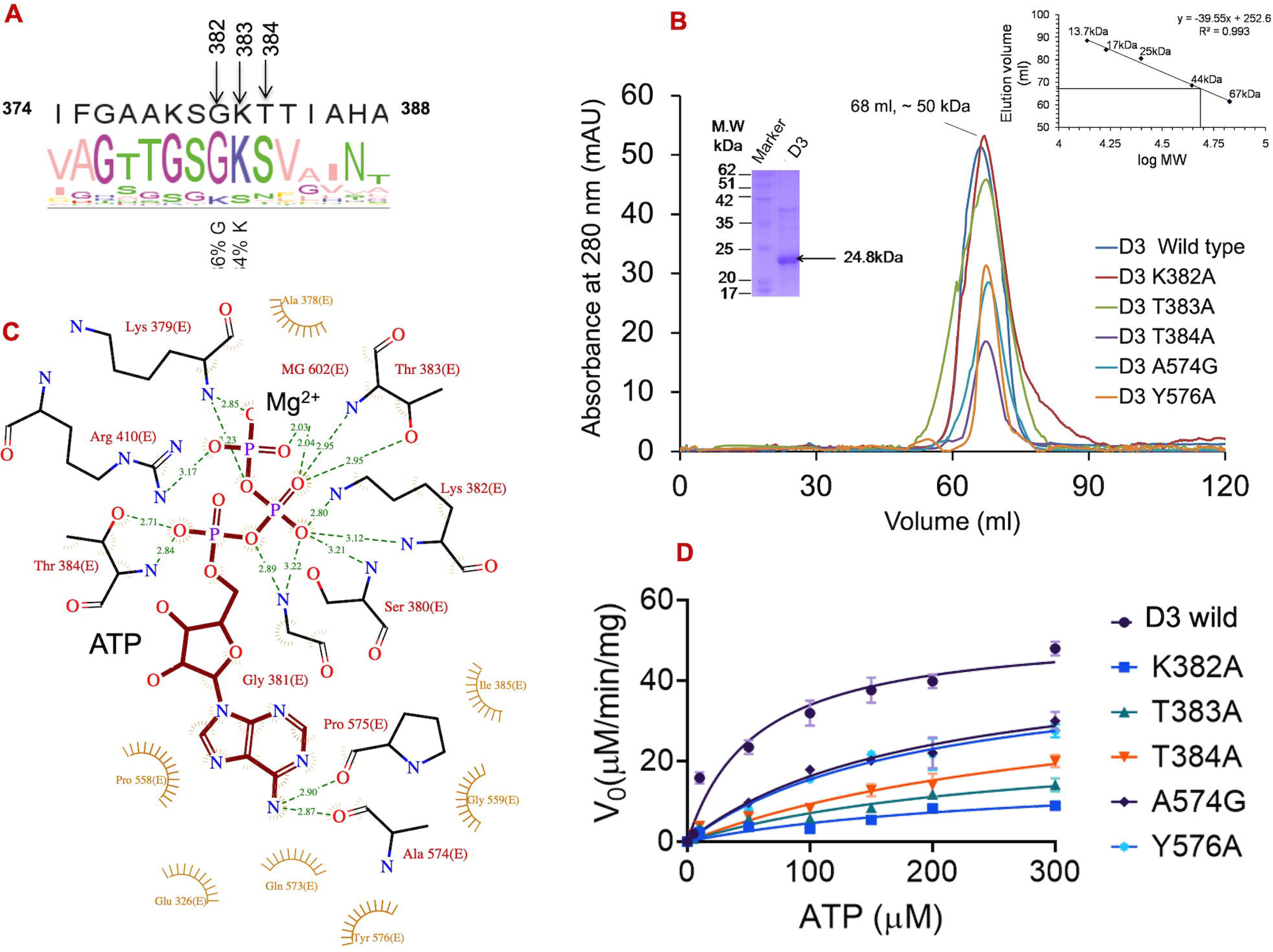
Purification and ATPase activity analysis of wild and mutant D3 domains. **A,** Web logo diagram of ATPase motif of D3 domain aligned with Ftsk/SpoEIII motif of 80 EccC sequences. **B,** Elution profile of wild type and five mutant D3 domains obtained from Superdex^TM^ 200 (16/60) column. The D3 domain eluted as dimer (based on molecular mass standard, *inset*) and protein appeared as single band on SDS-PAGE (*inset*) **C,** LigPlot analysis showing the interactions between ATP+Mg^2+^ and D3 domain. The ATP is shown in maroon, Mg^2+^ in green and D3 residues in red. **D,** Michaelis-Menten plots showing the ATP hydrolysis profile of wild D3 domain and its K382A, T383A, T384A, A574A, Y576A mutants.

The ATPase activity of D3 domain was determined using calorimetric assay and following parameters (*K_m_* ∼ 0.06±0.03 mM, catalytic efficiency (*k_cat_* /*K_m_*_)_ ∼ 0.030±0.008 μM^-1^ min^-1^ and *V_max_* ∼ 52.89 M/min) were obtained **(Fig. 4D).** Residues involved in ATP and Mg^2+^ binding were identified from *M. tuberculosis* ATPase3 domain+peptide+ATP+Mg^2+^ crystal structure (PDB-6J19). LigPlot analysis **(Fig. 4C)** showed that K382, T383, T384 residues of D3 domain bind to PO_4_ moiety and A574 and Y576 residues to Adenine ring of ATP.

To understand the roles of these residues in ATPase activity, K382A, T383A, T384A, A574G and Y576A mutants were generated using site directed mutagenesis **(Fig. 4C)** and ATPase assay was performed on all 5 mutants. **Table 1** showed the kinetic parameters obtained for 5 D3 mutants. These data showed that K382A mutation leads to (∼ 93.4 % decrease in catalytic efficiency, 80% decrease in V_max_). The T383A mutation leads to (∼ 89.7 % decrease in catalytic efficiency, ∼ 50% decrease in V_max_). The T384A mutation leads to (∼ 87.3% decrease in catalytic efficiency and ∼ 22% decrease in V_max._). The T574A mutation leads to (∼73.5% decrease in catalytic efficiency and ∼ 12% decrease in V_max_) and Y576A mutation leads to (∼75.2% decrease in catalytic efficiency and ∼ 15% decrease in V_max_), compared to wild type D3 domain. These data showed that K382A mutation is the most critical and leads to 93.4% decrease in catalytic efficiency of the enzyme. The wild type and five D3 mutants were expressed and purified under similar condition and showed no abnormal migration on size exclusion column **(Fig. 4B).** In all D3 mutants, difference in ATPase activities were not due to defect in folding or aggregation of proteins.

We have compared the kinetic parameters of D2 and D3 domains with full length EccCb1enzyme **(Table 1).** These data showed that D2 domain exhibited ∼ 6.9-fold higher V_max_ and ∼ 3.4 fold higher catalytic efficiency. The D3 domain exhibited ∼ 5.6 fold higher V_max_ and ∼ 1.7 fold higher catalytic efficiency compared to wild type EccCb1 enzyme.

### Modeling of EccCb1+ATP+Mg^2+^+EsxAB complex and EsxAB substrate binding analysis

The EccCb1+ATP+Mg^2+^ model was obtained using PDB-4N1A [10] and PDB-6J19 [16] structures as input templates *(as described in Materials and Methods section)*. The topology diagram of EccCb1 (**Fig. 5A)** showed the arrangement of α-helices (red) and β-sheets (pink) in EccCb1enzyme and numbered accordingly. The EccCb1+ ATP+Mg^2+^+EsxAB model contains total 28-580 residues, in which D2 domain contains (35-316 residues) and D3 domain contains (349-580 residues) and connected by linker of 317-348 residues (**Fig. 5B**). The D2 and D3 domains contain core 8 stranded central β-sheets, decorated by α-helices on either side of β-sheets. The EsxAB substrate binds to specialized groove at C-terminal region of D3 domain **(Fig. 5B).** PDBsum server [22] analysis showed the single hydrogen bond and multiple van der waals interactions involved in EsxB substrate binding (**Fig. 5C**). Binding analysis between EccCb1 and EsxAB substrate was performed using bilayer interferometry, which yielded the *K_D_* ∼34.4 nM **(Fig. 5D).** In presence of 0.02 mM ATP in buffer, the *K_D_* ∼ 11.4 nM was observed **(Fig. 5E).** These data showed that EccCb1 binds strongly to EsxAB substrate, and its affinity enhanced ∼ 3-fold towards substrate in presence of ATP, indicating ATPase energy involvement in substrate translocation.

**Fig. 5.**
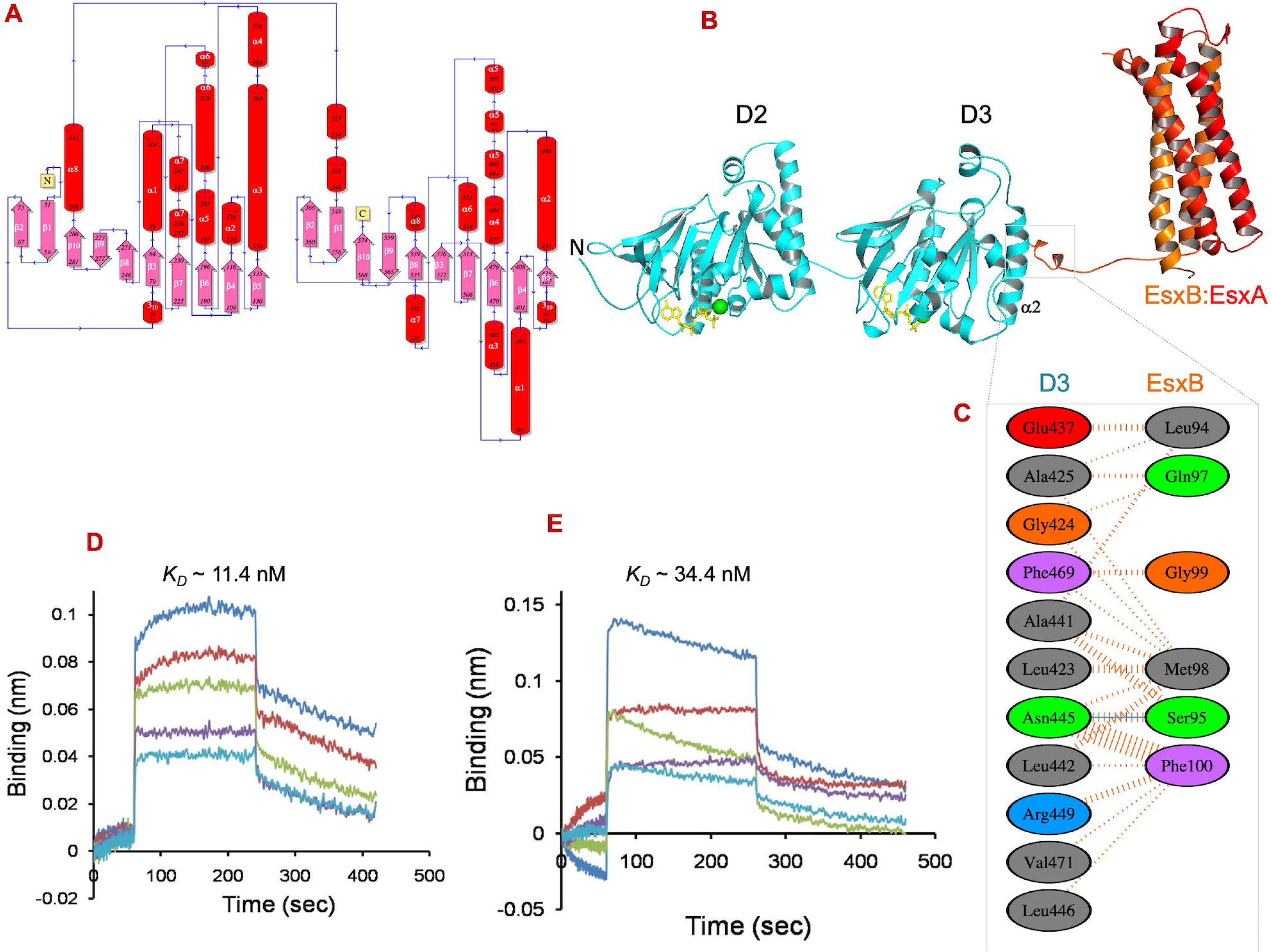
EsxAB substrate binding analysis using wild type and ATP bound EccCb1 enzyme. **A,** Electrostatic surface diagram of EccCb1 hexamer contoured at ±10kT. The EsxA (cyan) and EsxB (orange) bind to partially positively charged groove at C-terminal of D3 domain. The central cavity of EccCb1 is highly negative. **B,** The EccCb1+ATP+Mg^2+^+ EsxAB complex model showing the D3 domain pocket involved in binding to export arm of EsxAB virulence protein. **C,** showing the hydrogen bonds and vander waals interactions between EccCb1 and EsxB virulence protein. **D,** Sensogram showing the interaction between EsxAB heterodimer and EccCb1 in absence of ATP. **E,** Sensogram showing the interaction between EsxAB heterodimer and EccCb1 in presence of 0.3 mM ATP. The *K_D_* represents the binding affinity, *K_on_* is the association constant and *K_off_* is the dissociation constant.

### EccCb1 sequence alignment and comparative structure analysis

Sequence alignment of EccCb1 (1-591 residues) with *T. curvata* EccC enzyme (285-861 residues, PDB-4N1A) showed 39.8% sequence identity and 20.3% identity with and *G. thermodentroficans* EccC enzyme (1-419 residues, PDB -4LYA). The P-loop residues of D2 and D3 domains (shown as #) were found quite conserved to *T. curvata* EccC and *G. thermodentroficans EccC* enzyme sequences (**Fig. 6A)** and with 80 heterologous EccC sequences **(Fig. S3).** Superposition of EccCb1 structure on *T. curvata EccC* structure (PDB-4N1A) has yielded RMSD ∼ 0.78 Å for 461 Cα atoms and RMSD ∼ 7.1 Å for 428 Cα atoms with *G. thermodentroficans* EccC (PDB-4LYA) enzyme. The core α-helices and β-sheets of EccCb1 superposed very well with both enzymes, except minor differences in loop regions **(Fig. 6B).** In EccCb1 enzyme, D2 domain (28-316 residues) showed xxx identity with D3 domain (349-580) and superposition both domains yielded RMSD ∼ 5.7 Å over 146 Cα atoms (**Fig. 6C**).

**Fig. 6.**
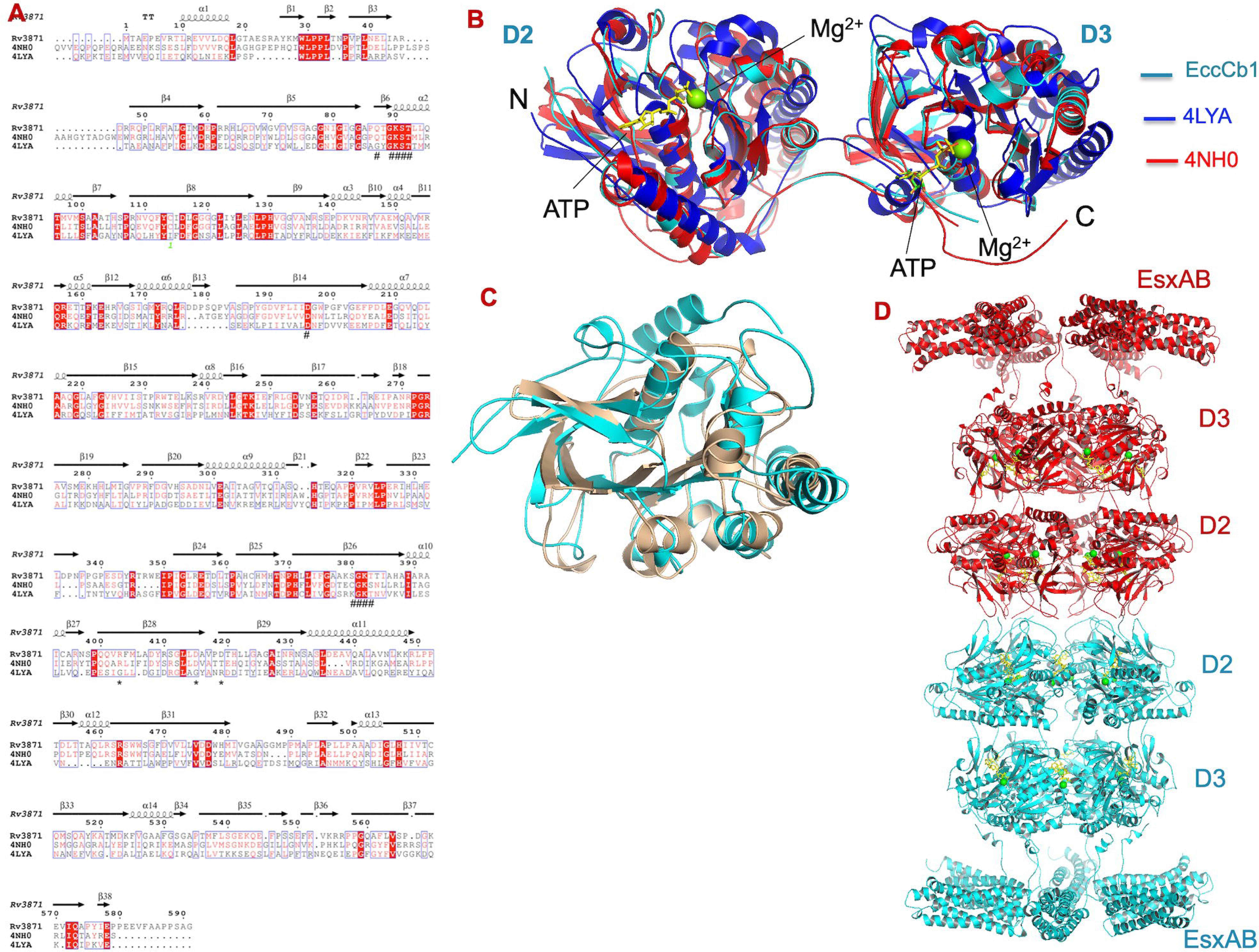
Sequence alignment, molecular modeling and comparative structure analysis of EccCb1 enzyme. **A,** Sequence alignment of EccCb1 (Cyam) with two structural homologs e. g. *T. curvata* EccC enzyme (PDB - 4NH0, red) and **D,** *EccC* enzyme (PDB- 4YVA, blue)) using MultiAln and Espript 5 programs. Semi conserved residues are shown in red letters and highly conserved in red shade. The EccCb1 residues involved in ATP+Mg^2+^ binding are shown in (#) above sequence alignment and EsxB binding in (*) below the sequence alignment. The β-strands are shown in black arrow and α-helices in spiral. **B,** Superposition of *T. curvata* EccC (PDB = 4NH0, cyan) and *D. EccC* (PDB- 4LYA, green) structures on EccCb1+ATP+Mg^2+^ structure (red) using PyMol program. **C,** Superposition of D2 domain on D3 domain. D2 domain is shown in orange and D3 domain in cyan. **D,** The EccCb1+ATP+Mg^2+^+EsxAB hexameric dimer viewed along helical axis. Each complex is shown with red and cyan colors respectively. ATP is shown in yellow and Mg^2+^ in green colors respectively.

The double hexameric EccCb1+EsxAB+ATP+Mg^2+^ complex model was obtained as shown in (*Materials and methods section*) (**Fig. 6D**). The double hexameric model showed the following dimensions, [21 (inner) x 94 Å (outer)] diameter for D2 ring and [21 (inner) x 102 Å (outer)] diameter for D3 ring and 80 Å height of the model **(Fig. 6D)**.

### Dynamics simulation analysis revealed the EccCb1 dynamics involved in EsxAB substrate and ATP+Mg^2+^ recognition

To study the EccCb1 dynamics involved in ATP+Mg^2+^ and EsxAB substrate recognition, 1 ns dynamics simulation was performed on EccCb1 dodecamer in three environments, (i) as apo enzyme (ii) complex with ATP+Mg^2+^ and (iii) complex with ATP+ Mg^2+^+EsxAB substrate **(Table 4).** The simulated EccCb1 models were superposed on starting structures and conformational changes observed during simulation were analyzed **(Fig. 7A-C).**

**Fig. 7.**
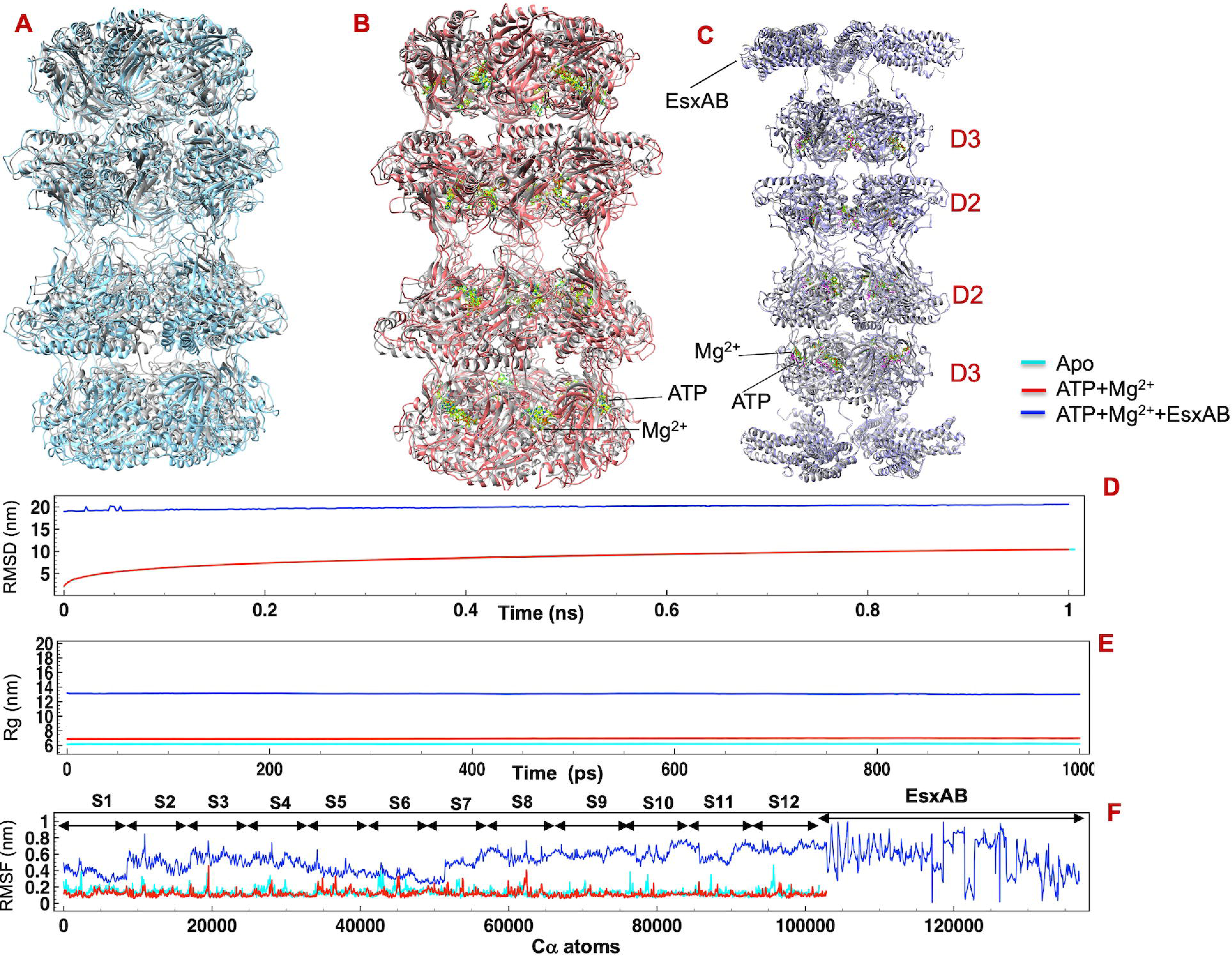
EccCb1 enzyme dynamics involved in ATP+Mg^2+^ and EsxAB substrate recognition. **A,** Superposition of the simulated apo EccCb1 dodecamer (cyan) on starting EccCb1 dodecamer (grey). **B,** Superposition of simulated EccCb1+ATP+Mg^2+^ dodecamer (red) on starting dodecamer (grey). **C,** Superposition of simulated EccCb1+ EsxAB +ATP+Mg^2+^ dodecamer (blue) on starting dodecamer (grey). **D,** RMSD of the EccCb1 models obtained during 1 ns dynamics simulation. **E,** Rg value of EccCb1 models obtained during 1 ns simulation. **F,** Plot showing the RMSF value of backbone Cα atoms of EccCb1 models.

**Table 3.** Binding analysis between EccCb1 and EsxAB substrate using Biolayer interferometry *(K_D_ is affinity constant, K_on_ is association constant, and K_off_ is dissociation constant)*.

| Immobilized | Ligand | ATP<br>(mM) | $K_{on}$<br>($10^5 \text{ M}^{-1} \text{ s}^{-1}$) | $K_{off}$<br>($10^{-3} \text{ s}^{-1}$) | $K_D$<br>(nm) |
| --- | --- | --- | --- | --- | --- |
| EccCb1 | EsxAB | 0.02 | 5.28±1.58 | 6.03±0.07 | 11.4±3.4 |
| EccCb1 | EsxAB | - | 2.22±0.43 | 7.16±0.12 | 34.3±6.7 |

**Table 4.** Details of various parameters used in dynamic simulation on EccCb1 dodecamer in three different environments.

| Models | Apo | ATP+Mg<sup>2+</sup> | ATP+Mg<sup>2+</sup>+EsxAB |
| --- | --- | --- | --- |
| Protein atoms | 50,814 | 50,814 | 50,814 |
| Water/Na<sup>+</sup> | 114,608/30 | 76,649/54 | 137,446/114 |
| Ligand atoms | - | 24 | 24/8640 |
| Minimization (Steep) | 50,000 | 50,000 | 50,0000 |
| NVT equilibration | 100 ps | 100 ps | 100 ps |
| NPT equilibration | 100 ps | 100 ps | 100 ps |
| Time steps (fs) | 2 | 2 | 2 |
| No. of steps | 500,000 | 500,000 | 500,000 |
| Simulation (ns) | 1 | 1 | 1 |

The residual flexibility of apo, ATP+Mg^2+^ bound and ATP+Mg^2+^+EsxAB bound EccCb1 dodecamer are shown in **Fig. 7D**. For apo EccCb1 dodecamer (cyan), the RMSD of backbone Cα atoms go from 0 to 5 nm in 200 ps and remained stable to 8 nm throughout 1 ns simulation and similar results were obtained for ATP+Mg^2+^ bound EccCb1 dodecamer (red). For EsxAB +ATP+Mg^2+^ bound complex (blue), the RMSD of backbone Cα atoms start from 19 nm and remained stable throughout 1 ns simulation. These data showed that backbone Cα fluctuations were maximum for wild type and ATP+Mg^2+^ bound EccCb1 dodecamer during 1 ns simulation. However, backbone Cα fluctuations reduced significantly upon EsxAB substrate binding to EccCb1 ATP+Mg^2+^ dodecamer.

The compactness (R_g_) and associated stability of apo, ATP+Mg^2+^ and ATP+Mg^2+^+EsxAB bound EccCb1 dodecamer are shown in **Fig. 7E**. The radius of gyration of EccCb1 models remain constant during 1 ns simulation. Following radius of gyration ∼ 6.2 nm for wild type, ∼ 7.0 nm for ATP+Mg^2+^ bound and ∼13 nm for EsxAB +ATP+Mg^2+^ bound EccCb1 dodecamer were observed during dynamics simulation. These data showed that compactness (stability) of EccCb1 dodecamer increased as a result of ATP+Mg^2+^ and EsxAB+ATP+Mg^2+^ binding to enzyme.

The RMSF of the backbone Cα atoms were calculated for EccCb1 models and examined the atomic fluctuations occurred during 1 ns simulation **(Fig. 7F).** Following RMSF values were obtained e. g. ∼ 0.2-0.4 nm for apo (cyan), ∼ 0.2-0.4 nm for ATP+Mg^2+^ bound (red) and ∼ 0.2-0.8 nm for EsxAB+ATP+ Mg^2+^ bound (blue) EccCb1 dodecamer.

Superposition of simulated EccCb1 dodecamer on starting EccCb1 dodecamer **(Fig. 7A)** have yielded following (i) RMSD ∼ 1.37 Å (6636 Cα atoms) for wild type (ii) ∼ 1.65 Å (6636 Cα atoms) for ATP+Mg^2+^ bound **(Fig. 7B)** and (iii) ∼ 0.46 Å for 6636 enzyme Cα atoms and ∼2.7 Å for 2184 ligand Cα atoms f in case of EsxAB+ATP+Mg^2+^ bound EccCb1 dodecamer **(Fig. 7C).** The secondary structures of EccCb1 dodecamer superposed well with starting structure in all models and large movement occurred in the loop region of EccCb1 dodecamer. Major changes were observed in orientations of EsxAB heterodimer during dynamics simulation.

## Discussion

In current study, we have analyzed the SAXS structure, ATPase activity and EsxAB substrate binding of EccCb1 enzyme. Key residues of D2 and D3 domains of EccCb1 enzyme involved in ATPase activity were identified and characterized using wild type and mutant enzymes. *De novo* low-resolution envelope of EccCb1 was determined using small angle X-ray scattering technique, which fitted very well with modeled double EccCb1+ATP+Mg^2+^ hexameric structure. The EsxAB substrate binds to EccCb1 dodecamer with nM affinity and a specific pocket at C-terminal region of D3 domain was involved in binding. The EccCb1 dynamics involved in ATP+Mg^2+^ and EsxAB substrate recognition was analyzed using dynamics simulation technique. Our structural and biochemical analysis on EccCb1will contribute significantly in development of antivirulence inhibitors, which will prevent virulence factor secretion by ESX-1 system.

The AAA+ ATPases enzymes usually oligomerize to remain in active form. The AAA+/RecA ATPases usually form active hexamer in solution, a common characteristic of this family of enzymes. The *G. thermodenitrificans* EssC oligomerizes in absence of EsxB substrate. However, *T. curvata* EccC enzyme multimerizes upon binding to EsxB substrate. In our case, EccCb1 eluted as oligomer from size exclusion column and appeared as double hexameric envelope in solution using small angle X-ray scattering data. The EccCb1 does not require EsxAB substrate for its multimerization, as observed for *T. curvata* EccC enzyme.

The full length EccCb1 at its D2 and D3 domains have exhibited ATPase activity. In D2 and D3 domains, residues involved in ATPase activity were identified and characterized using wild type and mutantD2 and D3 domains. The K90A mutation in walker A motif of D2 domain and K382A mutation in walker A motif of D3 domain leads to ∼ 94% decrease in enzyme activity. In *T. curvata* EccC enzyme, T92A mutation in walker A motif of D2 domain and T383A mutation in D3 domain also lead to ∼ 90% decrease in enzyme activity,

The D2 and D3 domains showed higher ATPase activity than full length of EccCb1 enzyme. Similar results were obtained for G. *thermodenitrificans* EssC enzyme, which hydrolyze ATP at much lower rate, compared to its D2 and D3 domains. In *T. curvata EccC* enzyme, both D2 and D3 domains bind to ATP+Mg^2+^ and showed no ATPase activity. In *G. dentrificans* EssC enzyme, D3 domain does not bind to ATP. In *Staphylococcus EssC* enzyme, the D3 domain lacks the ATP binding motif, as its P-loop is converted into helix and occluded the ATP binding.

The EsxAB binds to EccCb1at specialized groove at C-terminal region of D3 domain, distant from ATPase pocket. ATP binding to EccCb1 enhanced its affinity towards EsxAB substrate, indicating ATPase energy involvement in EsxAB substrate translocation. In *T. curvata* EccC enzyme, the EsxB substrate also binds to D3 domain, which leads to enzyme multimerization. In VirD4 ATPase enzyme [23], the C-terminal region of substrate binds to enzyme before its secretion. In *G. dentrificans* EssC enzyme, the EsxB substrate binds to D3 domain, which induces its multimerization. In *T. curvata* EccC enzyme, ATP binding does not show any effect on EsxB substrate binding.

Low resolution envelope of EccCb1 was obtained using SAXS data, which fitted well with double hexameric EccCb1structure **(Fig. 5D).** The double hexameric structures were also obtained in other AAA+ATPases enzymes e. g. PatFtsk docecamer [24], archael MCM dodecamers [25] and SV40-L tag protein [26]. It appears that AAA+ATPases enzymes have propensity to form higher ordered hexamer upon oligomerization and may form a substrate transport channel powered by its ATPase energy.

In EccCb1+EsxAB +ATP+Mg^2+^ dodecamer, twelve D2 and D3 domains contain specific pockets and bind to ATP+Mg^2+^ binding, however specific groove involved in EsxAB substrate binding was observed in D3 domain only. Comparative structure analysis of EccCb1+ATP+Mg^2+^ monomer with two homologous *T. curvata* EccC and *G. dentrifican* EssC enzymes showed the overall conserved structure between enzymes and highly conserved P-loop involved in ATPase activity.

Dynamics simulation analysis on apo, ATP+Mg^2+^ bound and EsxAB +ATP+Mg^2+^ bound EccCb1 dodecamer showed the enzyme dynamics involved in ATP+Mg^2+^ and EsxAB substrate recognition. The ATP+Mg^2+^ binding enhanced the overall stability of EccCb1 dodecamer, as ATPase site residues showed lower RMSF values compared to apo enzyme. The EsxAB substrate binding further stabilized the substrate-binding groove ofD3 domain, as low RMSF values were observed for substrate binding residues. Our *in-silico* studies have rationalized the biochemical data observed for EccCb1 ATPase activity and EsxAB substrate binding. In summary, our structural and biochemical data on EccCb1 will provide new avenues for the development of antivirulence inhibitors to block virulence factor -1 system.

## Materials and Methods

### Expression and purification

#### Preparation of full length EccCb1 enzyme

Full-length EccCb1 gene (encoding Met1-Glu591 residues) was amplified using polymerase chain reaction using *M. tuberculosis H37Rv* genomic DNA. The amplified gene was cloned into *pMAL-p2X* vector using *BamH1* and *HindIII* restriction sites. The resulting *pMAL-EccCb1* plasmid contains the genes for maltose binding protein and factor Xa enzyme at N-terminus of EccCb1 gene. The EccCb1 plasmid was verified by gene sequencing and restriction-digestion analysis. The *pMAL-EccCb1* plasmid was transformed in *E. coli. ER2508* cells and single colony was grown in terrific broth media (Ampicillin ∼ 100 µg/ml) at 37°C, till OD_600_ ∼ 0.6-0.7. The cell culture was induced with 0.5 mM IPTG at 15 °C and grew the cells for another 18 hr. The MBP-EccCb1 fusion protein was overexpressed in soluble fraction of cell.

The cells were centrifuged at 8000 rpm for 3 min at 4 °C and suspended in lysis buffer-A (25mM Tris-HCl pH 7.5, 500 mM NaCl, 1 mM EDTA, 15%(v/v) Glycerol, 10mM Arginine, 4mM β-mercaptoethanol, 1mM Phenylmethylsulfonyl fluoride, 2 mM Benzamidine hexachloride, 0.3 mM ATP, 0.2 mg/ml Lysozyme). The cell lysate was homogenized, sonicated for 4 min and subjected to 25,000 x g centrifugation for 30 min to collect the supernatant. The supernatant was mixed with amylose resin (*from NEB*), pre-equilibrated with buffer-B (Buffer-A with no lysozyme) and loaded in empty column. The column was washed with buffer-B + 0.3 mM Maltose. The MBP-EccCb1 fusion protein was eluted in buffer-B +5mM Maltose.

For MBP tag cleavage, MBP-EccCb1 fusion protein was mixed with Factor Xa enzyme at [1000:1 (w/w) ratio] and dialyzed in buffer (20mM Tris-HCl pH 8.0, 100 mM NaCl, 2 mM CaCl_2_) for 12 hr at 4°C. The cleavage reaction was stopped with 2mM PMSF and reaction was analyzed on 8% SDS-PAGE. The reaction mixture was loaded on amylose column and cleaved EccCb1 was eluted from column. The cleaved EccCb1 was loaded on Superdex 200^TM^ (16/60) column (*from GE Healthcare Ltd*), preequilibrated with buffer (25 mM Tris-HCl pH 7.5, 300 mM NaCl, 15%(v/v) Glycerol, 10 mM Arginine, 10 mM MgCl_2,_ mM ATP, 4 mM β-mercaptoethanol). The EccCb1 fractions were pooled and concentrated using 30 kDa cutoff ultracentrifugal device (*Millipore*). The protein concentration was determined using Bradford assay and purity was analyzed on 8% SDS-PAGE.

#### Preparation of wild type and mutant D2 domains

The D2 domain gene (encoding 35-316 residues) was amplified using polymerase chain reaction using *M. tuberculosis H37Rv* genomic DNA. The amplified D2 domain gene was cloned into *pET23a* (+) expression vector using Nde1 and XhoI restriction sites. The *pET23a*-D2 domain gene plasmid was verified by gene sequencing and restriction-digestion analysis. The D2 plasmid was transformed in *E. coli. BL21(DE3)Plys* cells (Ampicillin ∼ 100 µg/ml) and single colony was grown in terrific broth media, till OD_600_ ∼ 0.6-0.7. The culture was induced with 1 mM IPTG at 25°C and grown for another 14 hr. The cells were harvested at 8000 rpm at 4 °C and suspended in lysis buffer-A (50mM Tris-HCl pH 7.5, 300mM NaCl, 15% Glycerol, 4 mM β-mercaptoethanol, 1 mM Phenylmethylsulfonyl fluoride, 2mM Benzamidinehexachloride, 10mM MgCl_2,_ 0.3mM ATP, 0.2 mg/ml Lysozyme and 10mM Arginine). The cell lysate was homogenized, sonicated and centrifuged at 25,000 x g to collect the supernatant.

The supernatant was loaded on Ni-NTA column (*Sigma*), preequilibrated with buffer-B (Buffer-A with no lysozyme and 0.3 mM ATP) at 4 °C. The column was washed with buffer-B+20mM imidazole and eluted the protein in buffer-B+250mM imidazole. The protein was loaded on Superdex200^TM^ (16/60) column (*GE Healthcare Ltd*), preequilibrated with buffer-B (50mM Tris-HCl pH 7.5, 15% (v/v) Glycerol, 10 mM Arginine, 300 mM NaCl, 10 mM MgCl_2,_ 4 mM β-mercaptoethanol, 0.3 mM ATP). The protein fractions were pooled and concentrated using 3kDa cutoff ultracentrifugal device (*Millipore*). The protein was analyzed on 12% SDS-PAGE and mass spectrometry. The protein concentration was determined using Bradford assay.

Site directed mutagenesis (*Stratagene Ltd*) was performed using *pET23a-D2* plasmid as template. Forward and reverse primers were designed to generate the Q87A, T92A, S91A and K90A mutants of D2 domain **(Table S1).** PCR reaction was performed, followed by DpnI digestion and transformed all plasmids in *E. coli* DH5α cells. Gene sequencing was performed on all four mutant plasmids to confirm the presence of correct mutation. All D2 mutant enzymes were overexpressed and purified using wild type D2 domain purification protocol.

#### Preparation of wild type and mutant D3 domains

The D3 domain gene encoding (349-580 residues) was amplified using *M. tuberculosis H37Rv* strain and cloned into *pET-SUMO* vector using TA cloning method. The resulting D3 gene plasmid was verified by gene sequencing. The D3 domain gene plasmid was transformed in *E. coli BL21(DE3)* cells (Kanamycin ∼ 150 μ colony till OD_600_ ∼ 0.6-0.7 at 37 °C. The cell culture was induced with 0.5 mM IPTG at 12°C and grew for another 20 hr. The cell culture was harvested at 14,000 x g and suspended the pellet in lysis buffer-A (50 mM Tris/HCl pH 7.5, 300 mM NaCl, 4 mM β mercaptoethanol, 15% Glycerol, 2 mM Benzamidine-hydrochloride, 1 mM Phenylmethylsulfonyl fluoride, 10 mM MgCl_2,_ 0.3 mM ATP, 0.2 mg/ml Lysozyme and 10 mM Arginine). The cell lysate was homogenized, sonicated and centrifuged at 25,000 x g at 4 °C.

The supernatant was loaded on Ni-NTA column, preequilibrated with buffer-B (Buffer-A with no Lysozyme) and washed two times with buffer-B+20mM imidazole. The protein was eluted in buffer-B + 250mM imidazole. The *pETSUMO*-D3 domain was loaded on Superdex 200^TM^ (16/60) column pre-equilibrated with buffer-B (50mM Tris-HCl pH 7.5, 10% Glycerol, 300mM NaCl, 10mM Arginine, 2mM β-mercaptoethanol, 0.3mM ATP and 10 mM MgCl_2_). The protein fractions were collected and analyzed on 12% SDS-PAGE.

For SUMO-tag cleavage, *pETSUMO*-D3 domain was mixed with SUMO protease in 1000:1 (w/w) ratio and dialyzed in buffer (50mM Tris-HCl pH 8.0, 0.2% Igepal and 1mM DTT) for 12 hr at 4°C. The reaction was stopped with 2 mM PMSF and loaded on Superdex200^TM^ HiLoad (16/60) column, preequilibrated with buffer-B. The peak fraction containing cleaved D3 domain were pooled, concentrated and analyzed on 12% SDS-PAGE.

Site directed mutagenesis (*Stratagene*) was performed using *pETSUMO-*D3 domain plasmid. Forward and reverse primers were designed to generate the five K382A, T383A, T384A, A574G and Y576A mutants of D3 domain plasmid **(Table S1).** The gene sequencing was performed on D3 mutant plasmids to confirm the correct mutation. All D3 domain mutants were overexpressed and purified by protocol used for wild type D3 purification.

#### Preparation of EsxAB heterodimer

The EsxA and EsxB proteins were overexpressed and purified as described in earlier paper [27]. In brief, the EsxA gene was cloned in *pET21a* vector and transformed in *E. coli BL21(DE3)* cells having antibiotics (Ampicillin ∼ 100 μg/ml). The EsxB gene was cloned in *pET28a* vector and transformed in *E. coli BL21(DE3)* cells having antibiotics (Kanamycin ∼ 50 μg/ml). Single colony was inoculated in 3 l luria bertani media with antibiotics and grew the cells at 37 °C till OD_600_ ∼ 0.6-0.7. The cells were induced with 1 mM IPTG and grew for another 4 hr. The cells were harvested at 8000 rpm for 3 min. The EsxA and EsxB proteins overexpressed in soluble fraction of cells. For complex preparation, EsxA and EsxB proteins were mixed in 1:1 molar ratio and analyzed on native-PAGE for complex formation and purified using Superdex 200^TM^(16/60) Hiprep column.

### ATPase activity

#### ATPase activity of full length EccCb1

For ATPase activity analysis, EccCb1 was dissolved in ATPase buffer containing (25mM Tris-HCl pH 7.5, 10mM MgCl_2_). 0.5 µCi of [®^32P^]ATP was added in ATPase buffer and incubated at 37 °C. 1 µl of reaction mixture was spotted after every 15 min on TLC plate. The 0.5 M formic acid and 0.5 M LiCl solution were used to develop TLC plate and dried at 37 °C. The TLC plate was exposed to Fuji film BAS-MS 2025 imaging plate for 12 hr and analyzed using Typhoon FLA 9500 *(GE Healthcare Ltd).* The background obtained from reaction buffer (having no EccCb1) was corrected from each measurement.

Colorimetric assay was performed to measure the EccCb1 ATPase activity using ATPase assay kit (*Innova bioscience*). The ATPase assay was performed at 37°C for 30 min with 1 μl of EccCb1 (2 μg) in 200 μl of buffer (20mM HEPES pH 7.5, 150mM NaCl, 5% Glycerol, 2mM β-mercaptoethanol, 10 mM Mg^2+^ and variable amount of ATP) in 96 well plate. The dye buffer containing (120mM malachite green, 0.06% polyvinylalcohol, 6mM ammonium heptamolybdate, 4.2% sodium citrate) was added in the reaction buffer. After incubating for 15 min, 10 μl solution from every reaction mixture was transferred in 96 well plates and absorbance at 630 nm was measured. Absorbance from reaction buffer (having no EccCb1) was subtracted from each data. Based on the absorbance from phosphate standard curve, the release of inorganic phosphate was measured. Each assay was performed three times and average ATPase activity was calculated. The kinetic parameters (*K_m_, V_max_* and *K_cat_*) were calculated using Prism 6.0.2 program.

ATPase activity of EccCb1 was measured using [□^32P^] ATP as substrate at different time intervals in presence of Mg^2+^, Mn^2+^, Ca^2+^, EDTA and high NaCl concentration. The EccCb1 was dialyzed in buffer 25mM Tris-HCl pH 7.5 for 12 hr at 4°C. The EccCb1 was added in reaction buffer (25 mM Tris-HCl pH7.5, 150 mM NaCl, 5% Glycerol, 2mM β-mercaptoethanol, 10mM MgCl_2_, 1mM ATP and 0.5 μCi of [□^32P^] ATP). The reaction mixture was kept at 37°C for 1 hr and 1 μl of solution was spotted on TLC plate, dried and developed using 0.5M LiCl and 1M formic acid. The plate was dried, kept in IP for 12 hr and scanned using Typhoon FLA 9500. Intensity of each spot was measured using Image J software [28]. Similar experiments were performed with 10mM Ca^2+^ and 10mM Mn^2+^ to examine their effects on EccCb1 ATPase activity.

#### ATPase activity of wild type and mutant D2 domains

ATPase activity of wild type and four D2 mutants were determined using calorimetric assay using ATPase kit obtained from *INNOVA Biosciences Ltd*. 1 μg of D2 domain was dissolved in 200 μl of reaction buffer (50 mM Tris-HCl pH 7.5, 15% Glycerol, 10 mM MgCl_2_, 150 mM NaCl, 10mM Arginine) and incubated 10 min at 25 °C using various concentrations of ATP. 50 µl of Malachite green reagent was mixed in reaction mixture and incubate for 30 min. The absorbance at 600 nm was measured and absorbance from reaction buffer (no D2 domain) was subtracted from each data. The released inorganic phosphate was estimated using standard absorbance curve based on KH_2_PO_4_ assay. All assays were performed three times and kinetic parameters (*K_m_, V_max_* and *K_cat_*) were calculated using Prism 6.0.2 program (*GraphPad Software*). Similar protocol was used for ATPase activity analysis of four D2 mutant enzymes.

#### ATPase activity of wild type and mutant D3 domains

Calorimetric assay was performed for ATPase activity analysis of wild type and five D3 mutants using ATPase kit from *INNOVA Biosciences Ltd*. Initially, D3 domain was dialyzed in buffer (25 mM Tris-HCl pH 7.5, 0.5 mM β-mercaptoethanol, 150 mM NaCl, 5% Glycerol). 1 μM of D3 domain was added in 200 μl of reaction mixture (50 mM Tris-HCl pH 7.5, 15% Glycerol, 10 mM MgCl_2_, 150 mM NaCl, 10mM Arginine) from ATPase kit. The reaction mixture was incubated 30 min at 20 °C and 50 µl of Pi color lock was added in each well. After 2 min, 20 μl of stabilizer was added in each well, incubated for another 30 min and measured the absorbance at 630 nm. All assays were performed three times and kinetic parameters (*K_m_, V_max_* and *K_cat_*) were calculated using Prism 6.0.2 program (*GraphPad Software*). Similar protocol was used for ATPase activity measurement of all five D3 mutants.

### Small angle X-ray scattering analysis

Small angle X-ray scattering data on EccCb1 was collected using 1D Dectris Mythen detector at 10 °C using 0.154 nm wavelength at SAXSpace equipment at IMTECH, India (Table 1). 40 μl of EccCb1 enzyme (∼ 5 mg/ml) in 50mM Tris-HCl pH 7.5, 4mM β mercaptoethanol, 300mM NaCl buffer was loaded on thermostated 1 mm quartz capillary and exposed 30 min. Three frames were collected and averaged the data for protein and buffer. SAXSTreat program was used to correct the beam position, to subtract the buffer contribution and desmearing of scattering profile. The SAXS intensity profile was obtained as I(q) ∼ q (momentum transfer vector). ATSAS suite of programs (version 3.0.1) [29] was used for all further data analysis.

The scattering profile of EccCb1 in solution was analyzed using Kratky plot (I(q) ∼ q^2^). Guinier analysis was performed to obtain the R_g_ (radius of gyration) of EccCb1 in solution. Using I(q) profile, P(r) curve and estimated parameters, the DAMMIF program of ATSAS suite [30] was used for *ab initio* shape determination using dummy atom model of EccCb1. DAMMIF program [30] generated the ten independent envelopes after aligning on inertial axes, comparison, clustering in family-based similarity and averaging. These *de novo* models were fitted with homology models of EccCb1 by PyMol based plugin SUPALM program [31].

### Biolayer interferometry analysis

The interaction of wild type Rv3871 and its virulence factor (export arm of EsxAB heterodimer) was carried out using Forte Bio Octet Red 96 instrument at 30°C with 1000 rpm agitation. 50μg/ ml of Rv3871 and its mutants were individually immobilized on AR2G sensor and the experiment was carried out in GREINER 96 Solid black flat-bottom well plate in running buffer containing 20mM HEPES pH 7.5 and 150mM NaCl. Five different concentrations of virulence factor were prepared (30μM to 15μM and 7.5μM to 3.8µM and 1.9 µM in running buffer. Monitoring the interactions was conducted as 60 sec for initial base line, 180 sec association, 180 sec dissociation and finally regeneration using glycine-HCl pH 3.0. Data analysis was done using Forte Bio Data Analysis 10 and 1:1 binding model was used for curve fitting and calculation of association constant, dissociation constant and equilibrium dissociation constant.

### Molecular modeling and dynamics simulation

#### Homology modeling

The X-ray coordinates of EsxAB heterodimer were obtained from PDB ∼ 1WA8 (Solution structure of the CFP-10-ESAT-6 complex, major virulence determinants of pathogenic Mycobacteria) [32]. The EccCb1 model (1-591 residues) was obtained from I-TASSER server [33] using default parameters. The LOMETS program [34] used the EccCb1 sequence to identify the best structure templates from PDB database. The best template with high Z-score (high threading alignment score) was obtained from SPICKER program [35] after structural assembly simulation, and predicted the best models based on the C-score. The I-TASSER server yielded the best EccCb1 model (23-591 residues) based *T. cvurvata* EccC crystal structure (PDB-4NH0).

The crystal structure of D3 domain (PDB-6J19, 320-580 residues) was inserted in to obtained EccCb1 model based on PDB-4NH0 as template. The ATP+Mg^2+^ ligands were placed in D2 domain (23-327 residues) using PDB-4NH0 structure and in D3 domain (320-580 residues) based on PDB-6J19 structure. Using EccCb1+peptide+ATP+Mg^2+^ crystal structure (PDB-6J19) as template, the EsxAB heterodimer was docked into EccCb1+ ATP+Mg^2+^ complex using COOT program [36], which yielded the entire EccCb1+EsxAB+ ATP+Mg^2+^ complex structure.

To improve EsxAB docking into D3 domain, induced fit docking module was used in GLIDE program [37] with 0.6-fold scaling on van der Waals radii and XP (extra precision scoring function) for docking analysis. All residues within 6 Å radii of EsxAB substrate were refined for prime site optimization. 10,000 cycles of energy minimization and 10,000 cycles of scoring were performed using Glidestone module of Glide program, which yielded the best EccCb1+EsxAB+ATP+Mg^2+^ complex model. The best complex model was selected based on energy, interaction, buried surface area and lowest IFD.

The best EccCb1+EsxAB+ATP+Mg^2+^ model was given as input in SymmDock server [38] and generated the EccCb1+EsxAB+ATP+Mg^2+^ hexamer using 6-fold symmetry as constraint. The best hexameric complex was selected based on scoring function and given as input in SymmDock server using 2 fold symmetry, which yielded the cylindrical dodecameric structure of EccCb1+EsxAB+ATP+Mg^2+^ complex. 50,000 steps of steepest descent minimization was performed on obtained EccCb1+EsxAB+ATP+Mg^2+^ dodecamer using GROMACS program [39]. Structural validation on obtained dodecameric complex PROCHECK program [40] program was structural validation yielded the good stereochemistry of obtained model.

The MultAln [41] and ESPript 3.0 [42] programs were used for multiple sequence alignment of EccCb1 model against *T. curvata* EccC amd *G. Dentrificans* EssC sequences, best structural homologs identified by I-TASSER server. Structural superposition of EccCb1 against both enzymes and fitting into observed SAXS envelope were performed using CHIMERA program [43].

#### Dynamics simulation on Apo, ATP+Mg^2+^ and ATP+Mg^2+^ +EsxAB bound EccCb1 dodecamer

The dynamics simulation on (i) apo EccCb1 dodecamer (ii) EccCb1+ATP+Mg^2+^ dodecamer and (iv) EccCb1+EsxAB+ATP+Mg^2+^ dodecamer were performed using in GROMACS 2020.1-MODIFIED program using CHARMM36 all atom force field [44] and TIP3P water molecules parameters [45]. Details of all parameters used in dynamics simulation of EccCb1 models are given in **Table 4**. The EccCb1 models were kept in cubic box extended by 1.0 nm from protein surface and solvated using explicit SPC water molecules. Counter ions were added to neutralize the charge of the system. Before dynamics simulation, 50,000 steps of steepest descents energy minimization were performed on EccCb1 models. The particle-mesh Ewald (PME) method was used for electrostatic interactions having following parameters e. g. grid spacing 1.0 Å, cutoff 10 Å, relative tolerance of 10^-6^ and 10 Å switching for Lennard-Jones interactions. LINCS algorithm [46] was used for constraining bonds involved in hydrogen bonding.

100 ps of NVT equilibration was performed after heating the whole system at 300K, followed by 100 ps NPT equilibration. Finally, dynamics simulation was performed under NPT condition with V- rescale (modified Berendsen thermostat) coupling using Gromacs 10.12 version program. The 1.0 coulomb cut off was applied and 2 fs time step was used in dynamics simulation analysis. After every 4 ps, the coordinates were saved for MD trajectory analysis. The pressure was maintained by Parrinello-rahman pressure coupling constant and coulomb cut off was applied during temperature coupling. The Plot2X program [47] for trajectories analysis, PyMoL [48] and CHIMERA programs for structure visualization.

## Acknowledgements

We acknowledge the financial support from Department of Science and Technology, New Delhi DST-project (No. DST-EMR/2016/000867) to conduct the current research work. Current work was also partially supported by UGC-SAP grant, UGC-resource networking grant, DST-PURSE grant. We acknowledge the UGC, India for Senior Research Fellowship support to Arkita. The staff from CIF facility of SLS, JNU is gratefully acknowledged. We acknowledge the help from Dr. Ashish and other staff at IMTECH facility for SAXS data collection.

## Author’s contributions

Arkita cloned, expressed, purified the wild type and mutant EccCb1 proteins, She performed all ATPase assays and binding analysis with native and mutant proteins. Ajay K. Saxena performed all small angle X-ray scattering structure analysis, molecular modeling, docking and dynamic simulation of all EccCb1 complexes. Ajay K. Saxena conceived, designed the strategies, analyzed the whole data and wrote the entire manuscript.

