## supplementary material for "Structural and biochemical analysis of ATPase activity and EsxAB substrate binding of *M. tuberculosis* EccCb1 enzyme"

**Table S1.** Primers used for gene cloning of wild type and mutant Rv3871 and D1 and D2 domains.

**Rv3871 gene cloning primer**

Native
F: 5´- GATCGGATCCATGACTGCTGAACCGGAAGTA -3´
R: 5´- CATGAAGCTTTTAACCGGCGCTTGGGGGTGC -3´

R403A mutant
O: 5´- CAGTCCCCAGCAGGTGCGGTTCATGCTCGCGGACT -3’
F: 5´- CAGTCCCCAGCAGGTGGCGTTCATGCTCGCGGACT -3´
R: 5´- AGTCCGCGAGCATGAACGCCACCTGCTGGGGACTG -3´

L414A mutant
O: 5´- CTACCGCTCGGGCCTGCTGGACGCGGTGCCGGACA -3´
F: 5´- CTACCGCTCGGGCCTGGCGGACGCGGTGCCGGACA -3´
R: 5 ´- TGTCCGGCACCGCGTCCGCCAGGCCCGAGCGGTAG -3´

D419A mutant
O: 5´- GCTGGACGCGGTGCCGGACACCCATCTGCTGGGCG -3´
F: 5´- GCTGGACGCGGTGCCGGCCACCCATCTGCTGGGCG -3´
R: 5´- CGCCCAGCAGATGGGTGGCCGGCACCGCGTCCAGC -3´

**D1 gene cloning primer**

Native
F: 5´- GATCCATATGAATCCGGTCCCGCTCAACGAGCTCAT -3´
R: 5´- CATGCTCGAGTTCGGTGTGCTGGGAAGCGATCT -3´

Q87A mutant
O: 5´- ATTGGGGGCGCACCTCAAACCGGGAAGTCGACGCTAC -3´
F: 5´- ATTGGGGGCGCACCTGCAACCGGGAAGTCGACGCTAC -3´
R: 5´- GTAGCGTCGACTTCCCGGTTGCAGGTGCGCCCCCAAT -3´

K90A mutant
O: 5´- GCGCACCTCAAACCGGGAAGTCGACGCTACTGCAGAC -3´
F: 5´- GCGCACCTCAAACCGGGGCGTCGACGCTACTGCAGAC -3´
R: 5´- GTCTGCAGTAGCGTCGACGCCCCGGTTTGAGGTGCGC -3´

S91A mutant
O: 5´- CACCTCAAACCGGGAAGTCGACGCTACTGCAGACGAT -3´
F: 5´- CACCTCAAACCGGGAAGGCGACGCTACTGCAGACGAT -3´

R: 5′- ATCGTCTGCAGTAGCGTCGCCTTCCCGGTTTGAGGTG -3’

T92A mutant

O: 5′- CTCAAACCGGGAAGTCGACGCTACTGCAGACGATGGT -3′

F: 5′- CTCAAACCGGGAAGTCGGCGCTACTGCAGACGATGGT -3′

R: 5′- ACCATCGTCTGCAGTAGCGCCGACTTCCCGGTTTGAG -3′

---

**D2 gene cloning primer**

Native

F: 5′- GATCCATATGGAGATTCCGATCGGCTTGCGCGA -3′

R: 5′- CATGCTCGAGTGGAGGCTCGATGTAGGGGGCCTG -3′

K382A mutant

F: 5`- TGCGGCCAAATCGGGCGCGACGACCATTGCCCACG 3`

R: 5`- CGTGGGCAATGGTCGTCGCGCCCGATTTGGCCGCA 3`

T383A mutant

F: 5`- GGCCAAATCGGGCAAGGCGACCATTGCCCACGCGA 3`

R: 5`- TCGCGTGGGCAATGGTCGCCTTGCCCGATTTGGCC 3`

T384A mutant

F: 5`- CAAATCGGGCAAGACGGCCATTGCCCACGCGATCG 3`

R: 5`- CGATCGCGTGGGCAATCGGCGTCTTGCCCGATTTG 3`

Y576A mutant

F: 5`- GTCATCCAGGCCCCCGCCATCGAGCCTCCA 3`

R: 5`- TGGAGGCTCGATGGGCGGGGCCTGGATGAC 3`

A574A mutant

F: 5′- GGCAAAGAGGTCATCCAGGGCCCCTACATCGAGCCTCCA 3′

R: 5′- TGGAGGCTCGATGTAGGGCCGCTGGATGACCTCTTTGCC 3′

**Fig. S1.** Sequence alignment of Ftsk/SpoEIII motif of D1 and D2 domains with 80 different EccC sequences.

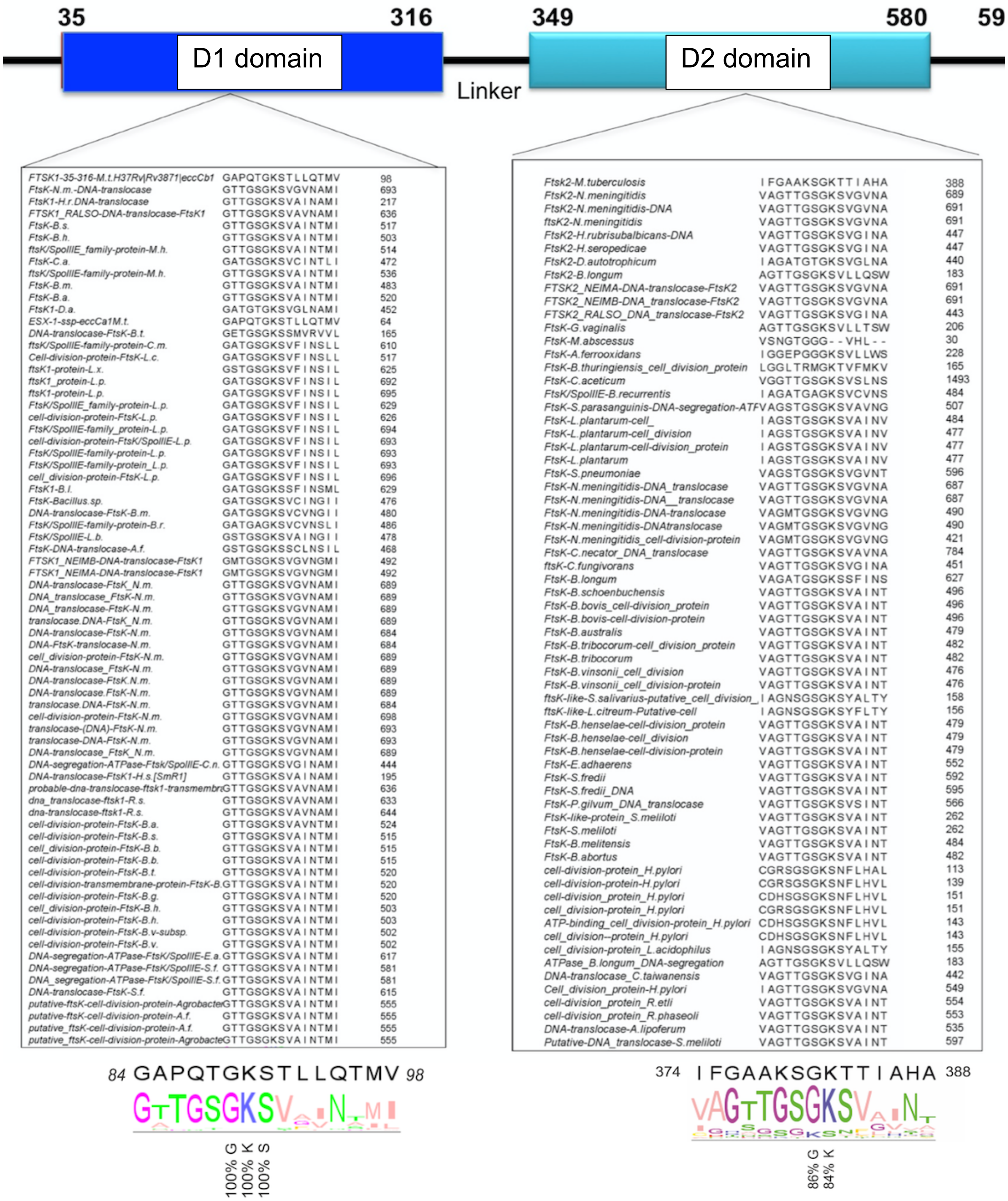
